# Distinct CASK domains control cardiac sodium channel membrane expression and focal adhesion anchoring

**DOI:** 10.1101/813030

**Authors:** Adeline Beuriot, Catherine A. Eichel, Gilles Dilanian, Florent Louault, Dario Melgari, Nicolas Doisne, Alain Coulombe, Stéphane N. Hatem, Elise Balse

## Abstract

Membrane-associated guanylate kinase (MAGUK) proteins function as adaptor proteins to mediate the recruitment and scaffolding of ion channels in the plasma membrane in various cell types. In the heart, the protein CASK (Calcium/CAlmodulin-dependent Serine protein Kinase) negatively regulates the main cardiac sodium channel, Na_V_1.5, which carries the sodium current (*I*_Na_) by preventing its anterograde trafficking. CASK is also a new member of the dystrophin-glycoprotein complex, and like syntrophin, binds to the C-terminal domain of the channel. Here we show that both L27B and GUK domains are required for the negative regulatory effect of CASK on *I*_Na_ and Na_V_1.5 surface expression and that the HOOK domain is essential for interaction with the cell adhesion dystrophin-glycoprotein complex. Thus, the multi-modular structure of CASK potentially provides the ability to control channel delivery at adhesion points in cardiomyocyte.

**SUMMARY:** Sequential functional domain deletion approach identifies three critical domains of CASK in cardiomyocytes. CASK binds the cell adhesion dystrophin-glycoprotein complex through HOOK domain and inhibits Na_V_1.5 channel membrane expression by impeding trafficking through L27B and GUK domains.

## INTRODUCTION

The nature of mechanisms regulating the delivery of newly addressed ion channels and their subsequent organization in the plasma membrane remains to be addressed in cardiomyocyte. Interestingly, members of the **M**embrane-**A**ssociated **GU**anylate **K**inase homologs (MAGUK proteins), a family of multi-domain membrane proteins, have emerged as central organizers of specialized plasma membrane domains in various cell types including cardiomyocytes. MAGUK proteins are able to interact with transmembrane proteins such as ion channels and receptors as well as with cytoplasmic proteins including scaffold, cytoskeleton and trafficking proteins, molecular motors, and signaling enzymes (Zhu et al., 2016). As such, MAGUK proteins have crucial roles in organization of cellular polarity, cell-cell adhesion, and dynamic regulation of large macromolecular complexes (Funke et al., 2005).

MAGUK proteins display a basic core structure comprising three main protein-protein interacting domains: PDZ (**p**ost-synaptic density 95, **d**isc **l**ar**g**e and **z**onula Occludens-1), SH3 (**s**rc **h**omology **3**), and GUK (**gu**anylate **k**inase) domains. In addition to this core architecture, the MAGUK protein CASK (**ca**lcium/calmodulin-dependent **s**erine protein **k**inase) comprises three additional domains that also serve as protein-protein interaction modules: an N-terminal CAMKII domain, a pair of L27 domains (L27A and L27B), and a HOOK domain preceding the C-terminal GUK domain. In neurons and epithelial cells, the role of each functional domain of CASK has been extensively investigated. For instance, the CAMKII domain interacts with Mint 1 and liprin-α, two proteins involved in vesicle exocytosis at presynaptic terminals (Butz et al., 1998; LaConte et al., 2016). The L27A (or L27N (terminal)) domain interacts with the MAGUK protein SAP97 in epithelial cells and targets the latter at the basolateral membrane (Lee et al., 2002). The L27B (or L27C (terminal)) binds to Veli, a multi-adaptator protein, and this association drives the basolateral distribution of Kir2.3 potassium channels (Alewine et al., 2007). The PDZ domain recognizes C-terminal sequences on target proteins, including transmembrane and cytoplasmic proteins. The SH3 domain binds GUK domains, enabling formation of CASK dimers (Nix et al., 2000), but also proline-rich motifs on target proteins, such as the C-terminal domain of calcium channel. This interaction involves Mint1 and helps channel trafficking to the plasma membrane and subsequent delivery of new cargo vesicles through Ca^2+^ influx (Maximov et al., 1999). The HOOK domain interacts with the cytoskeleton protein 4.1 and with adhesion molecules such as neurexin and syndecan-2 (Biederer and Sudhof, 2001; Chao et al., 2008). The GUK domain interacts with transcription factors to regulate the expression of genes containing T- or E-box consensus sequences (Hsueh et al., 2000; Wang et al., 2004; Ojeh et al., 2008). In addition, the GUK domain mediates the interaction between the presynaptic protein rabphilin3a and neurexin (Zhang et al., 2001).

Cardiomyocytes are structurally and functionally polarized cells (Balse et al., 2012). Intercalated discs, located at cell-cell endings, are responsible for electro-mechanical coupling and preferential propagation of the action potential. At the lateral membrane, two specialized domains are found: T-tubules, locus of the excitation-contraction coupling, and costameres which correspond to focal adhesions in non-muscle cells. We previously reported that in contrast to other MAGUK proteins identified in the myocardium, CASK is restricted to costameres, where it associates with the dystrophin-syntrophin complex. Using sh-RNA and overexpression approaches, we showed that CASK regulates the surface expression of the main cardiac sodium channel Na_V_1.5 in cardiomyocytes, and the corresponding sodium current *I*_Na_ by impeding Na_V_1.5 anterograde trafficking (Eichel et al., 2016). However, the mechanisms involved in these processes have not been fully explored.

Here we investigated the role of the different CASK domains in organizing and regulating the Na_V_1.5/dystrophin-glycoprotein macromolecular complex using CASK adenoviral constructs with sequential functional domain deletion: CASK^ΔCAMKII^, CASK^ΔL27A^, CASK^ΔL27B^, CASK^ΔPDZ^, CASK^ΔSH3^, CASK^ΔHOOK^ and CASK^ΔGUK^. Using a combination of whole-cell patch-clamp, total internal reflection fluorescence (TIRF) microscopy and biochemistry, we showed that L27B and GUK domains are involved in CASK negative regulatory effect on *I*_Na_ and Na_V_1.5 surface expression in cardiomyocytes. Using co-immunoprecipitation experiments, we showed that the HOOK domain of CASK is responsible for interactions with the dystrophin-glycoprotein complex. This study provides new insights into how the multi-modular structure of CASK makes the link between Na_V_1.5 functional expression at the membrane and the focal adhesion sites composed of the dystrophin-glycoprotein complex in cardiomyocyte.

## RESULTS and DISCUSSION

### Design and expression of truncated CASK adenoviral constructs in cardiomyocytes

We designed one full-length CASK protein corresponding to the 100 kDa cardiac isoform (CASK^WT^) and seven domain-specific deletion versions in which one single domain was truncated at a time (CASK^ΔX^). Deleted CASK proteins and full-length CASK were fused to a C-terminal GFP reporter (Figure 1A). GFP alone serves as a control condition. All constructs were sub-cloned into adenoviral vectors to enable overexpression in adult rat cardiomyocytes. Figure 1A shows the expected molecular weights of the various CASK isoforms. Western blot experiments performed on lysates obtained from cultured ventricular adult rat cardiomyocytes three days after adenoviral infection confirmed CASK protein overexpression at anticipated molecular weights (Figure 1B). A range of 100 to 140 kDa proteins corresponding to the full-length protein or truncated forms of CASK plus GFP was observed at significantly higher levels for CASK^WT^ and CASK^ΔX^ than the GFP control expressing only endogenous CASK (Figure 1B). RT-qPCR experiments performed on total mRNA isolated from cardiomyocytes three days after adenoviral infections demonstrated an average 6-fold increase in CASK mRNA levels for all eight constructs compared to the GFP control. Both CASK^WT^ and CASK^ΔX^ constructs had equivalent mRNA expression levels (P<0.001), supporting that all constructs had similar transduction efficacy in cardiomyocytes (Figure 1C).

**Figure 1.**
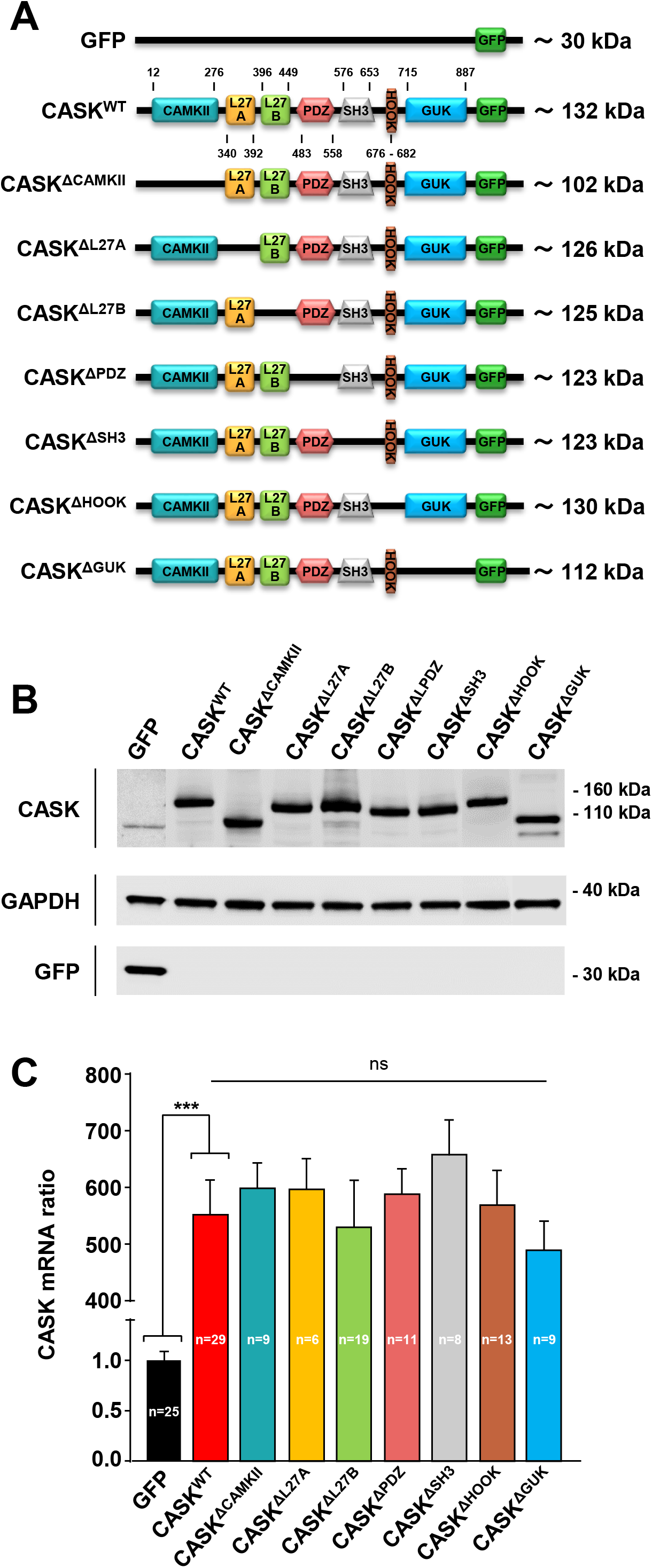
Validation of CASK adenoviral constructs in cardiomyocytes. **(A)** Schematic representation of CASK structure, domain-specific deletions, and expected molecular weights. **(B)** Representative western blot showing CASK expression in cultured rat cardiomyocytes transduced with each truncated CASK construct. CASK^WT^ fused to GFP corresponds to a ~140 kDa band. GAPDH served as a loading control. **(C)** RT-qPCR histograms of CASK mRNA expression levels in cardiomyocytes transduced with GFP, CASK^WT^, or CASK^ΔX^, where ΔX corresponds to any deletion. CASK mRNA levels in cardiac cells is normalized to endogenous CASK mRNA levels under control condition (GFP). Legend: ns, not significant; *** P<0.001; n=number of individual cell cultures; N=3-7 western blot replicates.

### Role of L27B and GUK CASK domains in sodium current regulation and Na_V_1.5 surface expression in cardiomyocyte

We previously showed that CASK acts as a negative regulatory partner of the cardiac sodium current *I*_Na_ by preventing anterograde trafficking and surface expression of Na_V_1.5 channels in the sarcolemma (Eichel et al., 2016). Here, we investigated the role of the different CASK domains in *I*_Na_ regulation and Na_V_1.5 surface expression.

First, whole-cell patch-clamp experiments were conducted in adult rat cardiomyocytes three days after transduction. For all constructs, i.e. CASK^WT^ and CASK^ΔX^, sodium current density was compared to GFP control only expressing endogenous CASK. *I*_Na_ was reduced by ~50% in CASK^WT^-transduced cardiomyocytes (P<0.001). Similarly, average current-voltage (I-V) plots from voltage clamp experiments showed significant decreases in *I*_Na_ densities at different tested voltages in CASK^ΔCAMKII^, CASK^ΔL27A^, CASK^ΔPDZ^, CASK^ΔSH3^, and CASK^ΔHOOK^-transduced cardiomyocytes (P<0.001). On the contrary, deletion of either L27B or GUK domains prevented such a reduction of *I*_Na_, and both CASK^ΔL27B^ and CASK^ΔGUK^ showed similar current densities to GFP control-transduced cardiomyocytes (Figure 1A-B). Cell capacitances were similar between the different conditions, implying that CASK does not significantly influence cell growth. Activation or steady-state inactivation properties were not changed between GFP, CASK^WT^, and CASK^ΔX^ (Figure 1C-D; Supplemental Table 1). These experiments demonstrate that CASK’s *I* Na inhibiting function is dependent on functional L27B and GUK domains.

**Figure 2.**
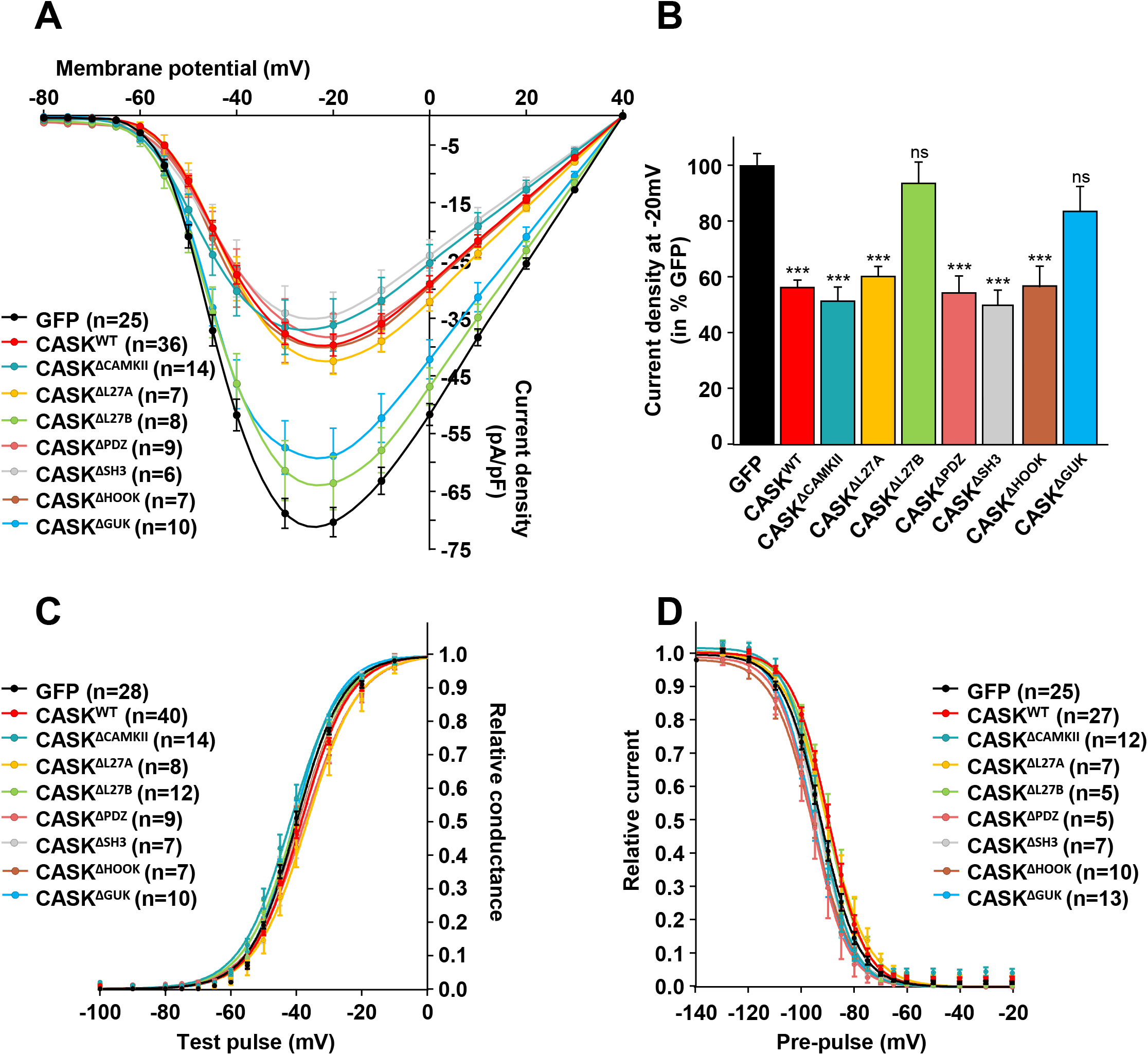
Both L27B and GUK CASK domains are implicated in the downregulation of the cardiac sodium current *I*_Na_ in cultured cardiomyocytes. **(A)** Current density-voltage relationships (25 mmol/L [Na^+^]_o_) of *I*_Na_ obtained from cardiomyocytes transduced with GFP control (black), CASK^WT^ (red), or deleted forms of CASK (CASK^ΔX^). **(B)** Histograms of *I*_Na_ current density recorded at −20mV and normalized to GFP control. **(C-D)** Voltage-dependent activation and steady-state inactivation curves from cardiomyocytes transduced with GFP control (black), CASK^WT^ (red), or truncated forms of CASK (CASK^ΔX^). Legend: ns, not significant; *** P<0.001; n=number of cells, N=3-16 independent cell cultures.

Second, total internal reflection fluorescence microscopy (TIRFm) was used to quantify the evanescent field fluorescence (EFF) intensity of the Na_V_1.5 signal in the membrane plane. Three days after transduction, cardiomyocytes seeded on TIRF-glass micro-dishes were fixed and stained with anti-Na_V_1.5 antibody. Cell boundaries were delineated using differential interference contrast (DIC) images and Na_V_1.5 EFF was quantified in critical angle in GFP control, CASK^WT^, CASK^ΔL27B^, and CASK^ΔGUK^ (Figure 3A). TIRF experiments showed a ~30% decrease in Na_V_1.5 surface staining in CASK^WT^ cardiomyocytes compared to GFP control (P<0.001) (Figure 3B). In contrast, the Na_V_1.5 EFF signal in either CASK^ΔL27B^- or CASK^ΔGUK^-transduced myocytes was of similar intensity to that measured in GFP control (Figure 3B).

**Figure 3.**
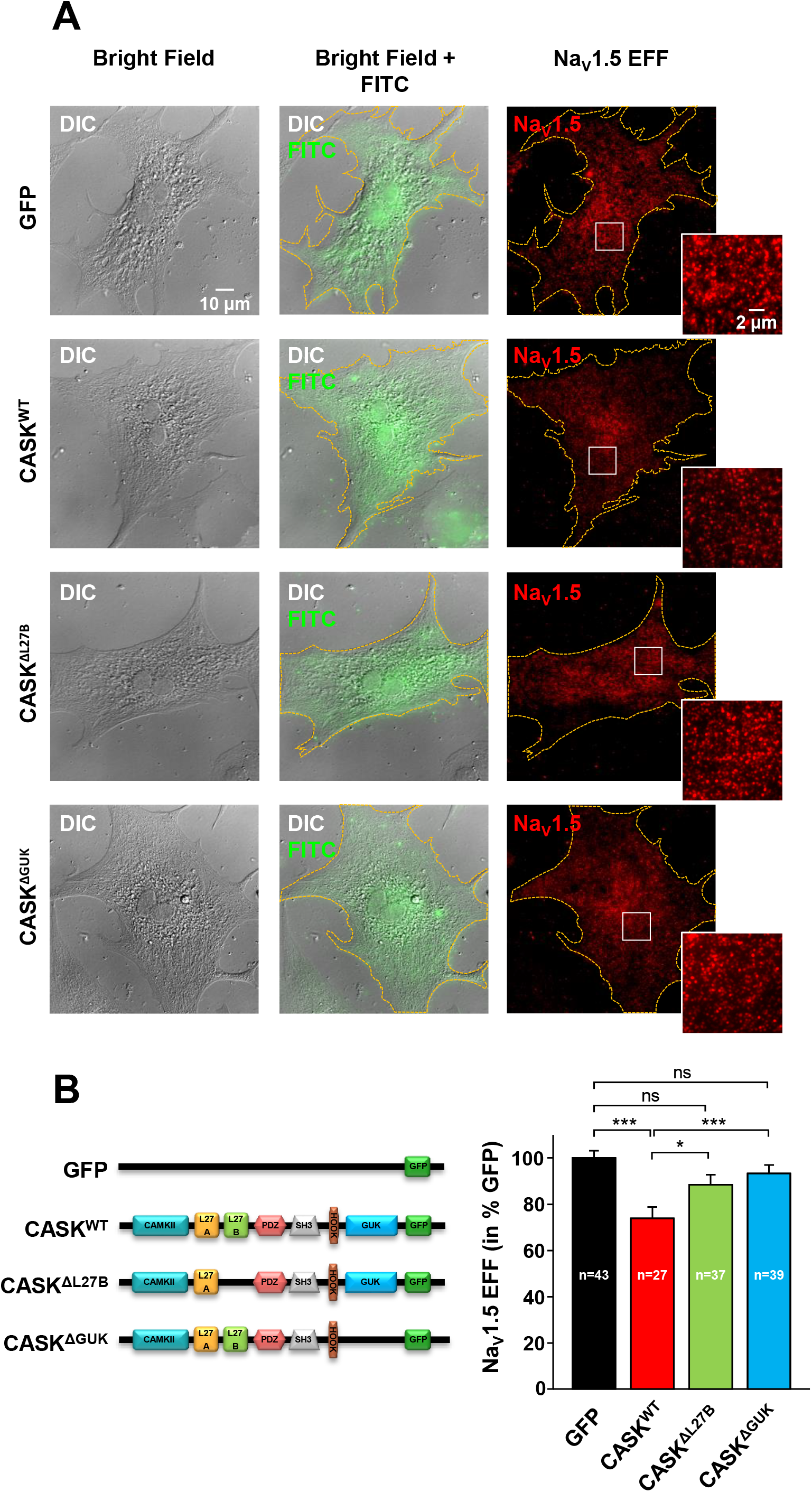
Deleting L27B or GUK CASK domains increases cardiac Na_V_1.5 channel surface expression. **(A)** Representatives differential interference contrast (DIC) and Total internal reflection fluorescence microscopy (TIRFm) images of both GFP and Na_V_1.5 signal taken from fixed cultured cardiomyocytes transduced with either GFP, CASK^WT^, CASK^ΔL27B^, or CASK^ΔGUK^. **(B)** Histograms of Na_V_1.5 evanescent field fluorescence (EFF) intensity in arbitrary units and normalized to GFP control. Legend: ns, not significant; * P < 0.05; *** P < 0.001; n=number of cells; N=4 independent cell cultures.

Third, RT-qPCR and Western blot experiments showed no changes in *scn5a* copy number and in global Na_V_1.5 protein expression between GFP control, CASK^WT^, and CASK^ΔX^ (Supplemental Figure 1), excluding the involvement of CASK in translational regulation of *SCN5A* gene encoding Na_V_1.5 channel.

Altogether, these results suggest that CASK negatively regulates *I*_Na_ by modulating Na_V_1.5 surface expression, independently of translational processes, through L27B and GUK domains. These results are particularly interesting considering the known functions of L27B and GUK domains of CASK in other cells types.

In neurons, correct synapse organization is ensured by the interaction of the extracellular domains of neurexin and neuroligin, respectively located in pre- and post-synaptic elements. Yeast two-hybrid screens identified CASK as an intracellular molecule interacting with neurexins (Hata et al., 1996). In the pre-synaptic terminal, a macromolecular complex formed by Mint1, a vesicular trafficking-related protein, Veli, a multi-adaptor protein, and CASK, has been characterized (Butz et al., 1998). CASK is the cornerstone of this complex as it binds Mint1 through its CAMKII domain and Veli through its L27B domain (Butz et al., 1998; Borg et al., 1999). Due to its interactions with both neurexin and vesicular trafficking-related proteins, this tripartite complex has been proposed to couple synaptic vesicle exocytosis to neuronal cell adhesion. In this line, the Mint1/Veli/CASK complex participates in anterograde trafficking of glutamate receptors along microtubules by bridging to the molecular motor kinesin, KIF-17 (Wong-Riley and Besharse, 2012). Other studies also reported interactions between the GUK domain of CASK and rabphilin-3a, a presynaptic protein involved in synaptic vesicle exocytosis. This suggests that the rabphilin-3a/CASK/neurexin complex could participate in the guidance of presynaptic vesicles towards exocytosis zones (Zhang et al., 2001). Consistent with these information, one can speculate that CASK, through its L27B and GUK domains, can regulate in a negative way the exocytosis of Na_V_1.5 channels in cardiomyocytes with the help of a not yet identified protein of the exocytosis pathway.

Additional proteins associate with the Mint1/Veli/SAP97/CASK complex, including SAP97, another MAGUK protein. Interestingly, the molecular arrangement of the complex affects glutamate receptors binding, trafficking and final targeting. For instance, differential sorting of glutamate receptor subtypes was shown to depend on SAP97/CASK interactions. When unbound from CASK, SAP97 preferentially associates with AMPARs, which are then trafficked to the plasma membrane through the classical secretory pathway. When bound to CASK, SAP97 associates with NMDARs which are then driven towards dendritic outposts instead of being retained in the endoplasmic reticulum (Jeyifous et al., 2009; Lin et al., 2013).

In this line, the existence of distinct Na_V_1.5 channel subpopulations has been proposed in cardiomyocyte relying on association with specific partners and specific localization in the sarcolemma (Shy et al., 2014). Na_V_1.5 channels are highly concentrated at the intercalated disc where they associate with gap junctional, desmosomal, actin cytoskeleton-binding, and MAGUK proteins. Na_V_1.5 channels at the lateral membrane show a lower density and interact with syntrophin and CASK (Balse and Eichel, 2018). The consequence of this differential distribution is to favor anisotropic (longitudinal > transversal) propagation of the action potential in the myocardium (Spach, 1999). Despite all identified partners exert regulatory function on sodium current, their precise role in trafficking, scaffolding and/or stabilization has not been investigated, except for CASK which prevent trafficking to the lateral membrane during early stage of trafficking (i.e. ER to Golgi) (Eichel et al., 2016). Therefore, one could speculate that interaction with specific partner(s) and/or competition between partners at early stages of trafficking could determine the sorting and final targeting of the channel in the sarcolemma. Although the sequence of interactions of the different CASK domains has not been completely characterized yet, our data demonstrates that both CASK L27B and GUK domains are mandatory in regulating the functional expression of Na_V_1.5 channel in cardiomyocyte sarcolemma, and strongly support our previous observation that CASK modulates anterograde Na_V_1.5 trafficking in cardiomyocytes (Eichel et al., 2016).

### Role of the CASK HOOK domain in scaffolding of macromolecular complex

Using immunohistochemistry and co-immunoprecipitation assays, we previously reported that CASK associates with dystrophin at the lateral membrane of cardiomyocytes. Immunostaining performed on dystrophin-deficient mdx mouse ventricular cryosections showed a loss of CASK staining, suggesting that CASK expression at the lateral membrane depends on dystrophin expression (Eichel et al., 2016). Here, we investigated whether a specific CASK domain was involved in this interaction.

Western blot experiments showed that CASK^WT^ or CASK^ΔX^ overexpression did not modify dystrophin expression levels compared to GFP control (Figure 4A-B). The association between CASK and dystrophin was again supported by co-immunoprecipitation experiments performed on cardiomyocyte lysates overexpressing CASK^WT^ (Figure 4C). Dystrophin was able to precipitate CASK with all truncated CASK^ΔX^ constructs except for CASK^ΔHOOK^ (Figure 4C). These results suggest that the ~10 amino acids HOOK domain of CASK is responsible for interactions with the costameric dystrophin-glycoprotein complex in cardiomyocytes. The fact that CASK^ΔHOOK^ does not rescue *I*_Na_ suggests that the HOOK domain is not involved in channel trafficking, but instead likely involved in organizing focal adhesions, notably costameric complexes.

**Figure 4.**
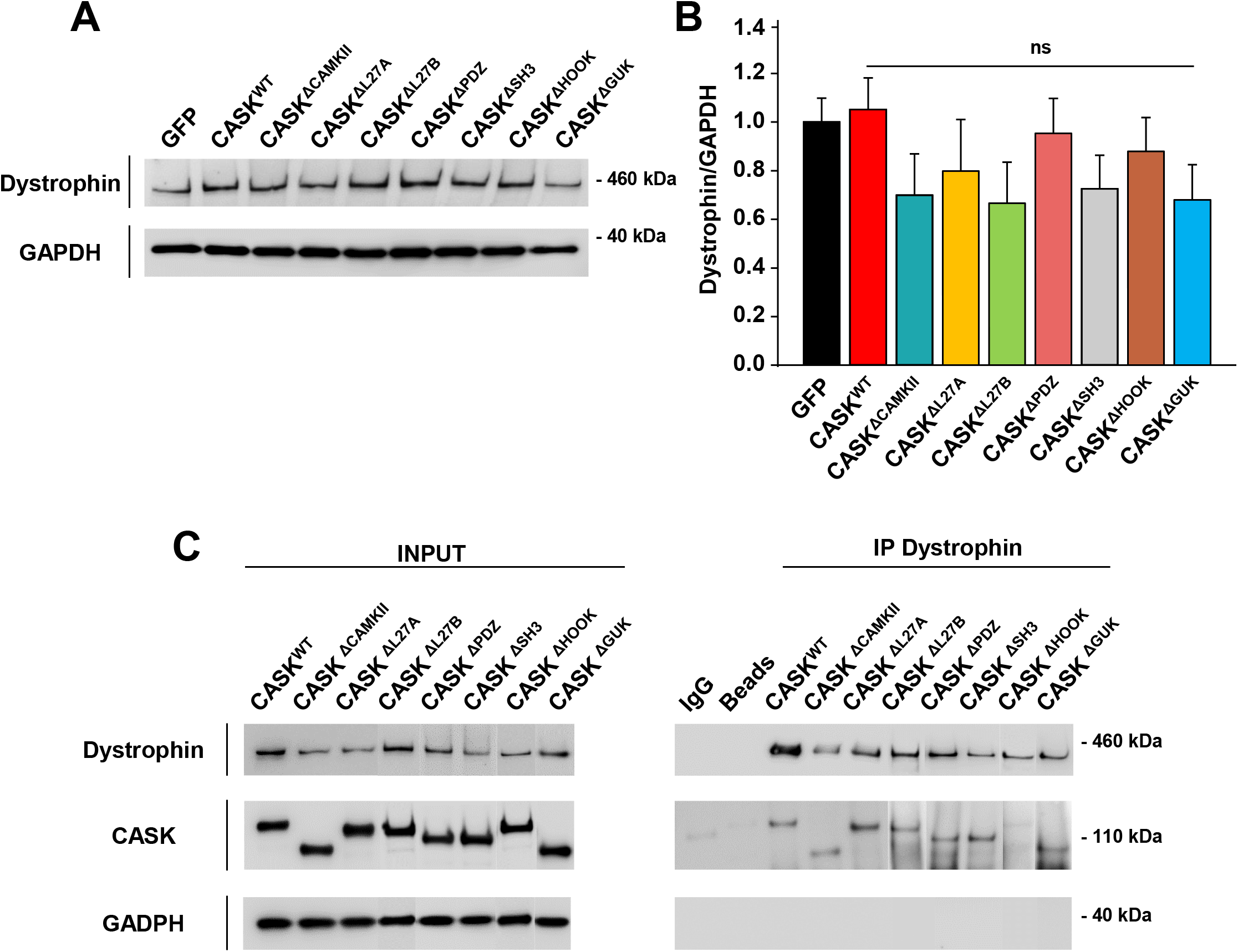
CASK interacts with dystrophin through its HOOK domain in cardiomyocytes. **(A)** Representative western blot of dystrophin expression after transduction with either GFP control, CASK^WT^, or CASK^ΔX^. **(B)** Corresponding histograms showing the expression level of dystrophin normalized to GAPDH in cardiomyocytes transduced with the different adenoviral constructs. Legend: ns, not significant; N=5 independent cell cultures. **(C)** Representative co-immunoprecipitation assays performed on cardiomyocyte lysates after transduction with CASK^WT^ or CASK^ΔX^ showing the loss of association with dystrophin for the CASK^ΔHOOK^ construct. N=3 independent cell cultures.

Costameres correspond to focal adhesions in cardiomyocytes and constitute physical links between sarcomeric z-lines, the sarcolemma, and the extracellular matrix. They guarantee the three-dimensional organization of the myocardium when sheets of myocytes slide against each other during each contraction. Interestingly, the HOOK domain of CASK corresponds to a protein 4.1 binding motif (Hanada et al., 2003). Protein 4.1 links spectrin and actin in the cytoskeleton and play critical role in mechanical stability and membrane protein organization (Baines et al., 2014). In epithelial cells, transmembrane syndecan is anchored to the actin cytoskeleton though interactions between CASK and protein 4.1 (Cohen et al., 1998). In neurons, intercellular junctions formed by neurexin and neuroligin at the synapse are also connected to the actin cytoskeleton through CASK and protein 4.1 interactions (Biederer and Sudhof, 2001). A large macromolecular complex formed of CASK, neurexin, Mint1, Veli, protein 4.1, spectrin, and actin, organizes and stabilizes the pre-synaptic element (Biederer and Sudhof, 2001). Despite all members of this complex possess PDZ domains, they are not involved in the formation of this large complex, rendering these domains free for other interactions (Butz et al., 1998). Indeed, in heart and skeletal muscle cells, the Mint1/Veli/SAP97/CASK complex was also found to be associated with ion channels, notably Kir2.x channels. Kir2.x channels also interact with components of the dystrophin-glycoprotein complex including syntrophin, dystrophin, and dystrobrevin at the neuromuscular junction, although with lower affinities (Leonoudakis et al., 2004).

In this line, using GST pull-down experiments we previously showed a direct interaction between purified CASK protein and the Na_V_1.5 C-terminal (Eichel et al., 2016). Na_V_1.5 channel was also shown to interact (directly or indirectly) with syntrophin (Gavillet et al., 2006). Despite the precise sequence of interaction and/or competition of the Na_V_1.5 channel with either syntrophin or CASK has not been investigated yet, the fact that CASK prevents anterograde trafficking of sodium channel could constitute a mechanism to maintain low levels of sodium channels at the lateral membrane. As already described for non-junctional Cx43/Na_V_1.5 channels in the perinexus at intercalated disc (Rhett et al., 2012), we hypothesize that the costameric macromolecular complex involving CASK constitutes a submembrane storage of non-functional sodium channels. We already showed that CASK expression in significantly reduced in remodeled/dilated atria, both in human and in rat model of heart failure (Eichel et al., 2016). Thus, changes in working conditions of the myocardium could affect Na_V_1.5 channel delivery from submembrane reservoirs to the lateral sarcolemma of cardiomyocytes and consequently alter anisotropic conduction.

## Conclusion

To the best of our knowledge, this is the first study to identify a sodium channel partner with the potential to couple ion channel trafficking to adhesion in specific area of cardiomyocytes, the focal adhesion of cardiomyocyte. Here, we show that CASK regulates anterograde Na_V_1.5 channel trafficking through L27B and GUK domains and interact with the dystrophin-glycoprotein complex, a mechano-transduction site, though HOOK domain. Due to its multi-modular structure, CASK could simultaneously interact with several targets in cardiomyocyte and potentially function as a cornerstone in locally directed exocytosis.

Mounting evidence indicates that distinct populations of Na_V_1.5 channels exist in cardiomyocytes, depending on their association with specific regulatory partners. However, the molecular mechanisms regulating temporal and spatial organization of Na_V_1.5 channels in the sarcolemma are still largely unknown. Therefore, exploring in real time the association of the sodium channel with its different partners and investigating further mechanisms leading to channel sorting and final targeting are now clearly warranted.

## MATERIAL and METHODS

### Animals

Studies were performed in adult male Wistar rats (~200 g, Janvier Labs, France). All animals were treated in accordance with French Institutional guidelines (Ministère de l’Agriculture, France, authorization 75-1090) and conformed to the Directive 2010/63/EU of the European Parliament for the use and care of animals.

### Isolation and culture of adult rat cardiomyocytes

Adult atrial or ventricular cardiomyocytes were obtained by enzymatic dissociation on a Langendorff column as previously described (Eichel et al., 2016). Briefly, rats were anesthetized with an i.p. injection of pentobarbital sodium (110 mg/kg) and heparin sodium (250 units/kg). The heart was then excised and cannulated via the aorta on a Langendorff apparatus and perfused in a constant flow rate (5mL/min). The perfusate consisted as an enzymatic solution containing in mmol/L: NaCl 100, KCl 4, MgCl2 1.7, glucose 10, NaHCO_3_ 5.5, KH_2_PO_4_ 1, HEPES 22, BDM 15 and 200UI/mL collagenase A (Roche Diagnostics) plus bovine serum albumin 7.5, pH 7.4 (NaOH) for ~20 min. Solution was equilibrate with 95% O2 and 5% CO_2_ and maintained at a temperature of 37°C. Atrial and ventricular tissues were separated and cut into small pieces and cells were gently dissociated with pipettes in Kraft-Brühe (KB) solution, in mmol/L: glutamate acid potassium salt monohydrate 70, KCl 25, taurine 20, KH_2_PO_4_ 10, MgCl_2_ 3, EGTA 0.5, glucose 10, and HEPES 10. Cardiomyocytes were plated on Petri dishes or Ibidi micro-dishes double-coated with Poly-D-lysine/laminin (Roche/Sigma). Cells were maintained under standard conditions (37°C, 5% CO2) in M199 medium (Gibco) supplemented with 1% penicillin/streptomycin (Gibco), 5% fetal bovine serum (Gibco) and 1‰ insulin/transferrin/selenium cocktail (Sigma). Isolated cells were used for immunofluorescence, biochemistry experiments or patch-clamp recordings after 3-5 days in culture.

### Adenoviruses

Full-length CASK (CASK^WT^) and truncated sequences (CASK^ΔCAMKII^, CASK^ΔL27A^, CASK^ΔL27B^, CASK^ΔPDZ^, CASK^ΔSH3^, CASK^ΔHOOK^, CASK^ΔGUK^) were designed and cloned into a pShuttle vector with CMV promotor. Expression cassettes were transferred into the adenovirus vector containing GFP reporter for selection of infected cells. All the production, amplification and purification process were conducted by a Vector Production Center (Gene Therapy Laboratory, INSERM U1089, IRT-1, Nantes, France). Rat adult cardiomyocytes were transduced 24h after isolation with 5.10^7 particles/ml of CASK^WT^, CASK^ΔX^ or empty vector (GFP) as negative control. Overexpression studies were performed 3 days post-transduction.

### Immunofluorescence and deconvolution microscopy

Cells fixation was performed with paraformaldehyde 4% (Sigma) for 10 min at room temperature (RT). For permeabilization, cells were incubated in PBS supplemented with 0.1% Triton X-100, 1% BSA, 10% sera for 1h at RT. Primary antibodies were applied over night at 4°C with blocking buffer (PBS containing 1% BSA, 3% sera, diluted antibodies according to the manufacturer’s instructions. Secondary antibodies (2.5 to 5 µg/mL) FITC (Abcam) or AlexaFluor594 (Life Technologies) and DAPI (0.2µg/µL Sigma) to detect nuclei were diluted in blocking buffer and added on samples for 1h incubation at RT.

### Total Internal Reflection Fluorescence microscopy (TIRFm)

Fluorescence intensity at surface membrane was visualized using the Olympus Cell^tirf^ system. Cells were placed on an Olympus IX81-ZDC2 inverted microscope and TIRF illumination was achieved with the motorized integrated TIRF illuminator combiner (MITICO-2L) and dual band optical filter (M2TIR488-561) using a 60X/1.49 APON OTIRF oil immersion objective, allowing GFP and RFP visualization. All image acquisitions, TIRF angle adjustment and some of the analysis were performed using the xcellence software (Olympus). Fluorescence analysis was performed using ImageJ software (NIH).

### Protein extraction

Cardiomyocytes expressing the appropriate constructs were lysed by scraping the cells into a lysis buffer, in mmol/L: Tris 50 (pH 7.5), NaCl 150, EDTA 2, 1% Triton X-100 and 1 ‰ protease cocktail inhibitor (Sigma Aldrich) and lysates were placed on a rotating 3D shaker for 1h at 4°C. After centrifugation, protein concentrations per supernatant were then determined by the BCA method (Pierce).

### Co-Immunoprecipitation assays

Cardiomyocytes were lysed as described above. Lysates were then incubated all night with antibodies (10µg) or IgG mouse (0.1µg/mL) at 4°C on a rotating 3D shaker. G-Sepharose beads (50µL for 2mg of total proteins, Sigma) were added for 1h before being washed with lysis buffer 3 times. Protein complexes bound to washed-beads were eluted with 2X sample buffer, followed by a denaturation step before separation by western blot.

### Western Blot analysis

NuPAGE^®^ Novex^®^ 10 % Bis-Tris or 3-8 % Tris acetate gels (Life Technologies) were used, depending on the size of expected proteins. Proteins were transferred onto nitrocellulose membrane (Biorad) and blocked with 5% nonfat dry milk for 60 min and probed overnight at 4°C with primary antibodies. Anti-rabbit or anti-mouse horseradish peroxidase-conjugated secondary antibodies (Cell Signaling) were used before revealing membrane with a chemiluminescent detection reagent ECL Prime (GE Healthcare). Images were acquired using a ImageQuant LAS4000 camera system (GE Healthcare) and then analyzed with the ImageJ software. The relevant spots were measured and normalized versus that of the corresponding spot in the control condition. These normalized densities were averaged from at least 3 independent experiments.

### Antibodies

Polyclonal rabbit antibody recognizing Na_V_1.5 was obtained from Alomone Labs (ASC-005, 8µg/mL). Monoclonal mouse anti-CASK used for biochemical assays were form Origene (Clone S56A-50, TA326522, 1.5-5µg/mL) and Merck Millipore (Clone 2G5.1, MABN1547, 0.6µg/mL). Monoclonal rabbit anti-GAPDH was purchased from Cell Signaling (Clone 14C10, 2118, 1:40000). Polyclonal rabbit anti-GFP used for immunochemistry (0.3µg/mL) and western blotting (2µg/mL) was obtained from CliniSciences (TP401). Monoclonal mouse anti-Dystrophin (Clone MANDYS8, D8168, 0.6µg/mL) used for biochemical assays were purchased from Sigma.

### Real-time reverse transcription polymerase chain reaction (RT-qPCR)

Total mRNA was extracted from cells in culture by using PureLink^®^ RNA Mini Kit (Life Technologies) and quantified by assessing optical density at 260 and 280 nm via a NanoDrop apparatus (Thermo Scientific). cDNA was synthesized using a mix of random primers, DNTP, Oligo DT12-18 follow by the addition of SuperscriptTM III reverse transcriptase (Life Technologies). Quantitative real-time PCR were carried out using SensiFAST™ SYBR No-ROX (2X, Bioline) on a LightCycler 480 System (Roche Diagnostics) according to the manufacturer’s specifications. All qPCR samples were run in duplicate and for each primer pair, a non-template control (NTC) was applied. Specific primers were designed to amplify *cask* (*cask*-F: CCAGGATCAGCATCTTCAC and *cask*-R: TGGCTCTCTGTACTGCATCG and *cask*-F: GTGAAGCCCCACTGTGTTCT and *cask*-R: TTCTGGGGCCATAAAGTGAG), *scn5a* (*snc5a*-F: TATGTCCTCAGCCCCTTCC and *scn5a*-R: GAACACGCAGTTGGTCAGG). All data were normalized to mRNA level of housekeeping genes *polr2a*, *rplp0* and *rpl32* (*polr2a*-F: CGTATCCGCATCATGAACAGTGA and *polr2a*-R: TCATCCATCTTATCCACCACCTCTT; *rplp0*-F: CGACCTGGAAGTCCAACTAC and *rplp0*-R: GTCTGCTCCCACAATGAAG; *rpl32*-F: CCAGAGGCATCGACAACA and *rpl32*-R: GCACTTCCAGCTCCTTGA) using the 2^−ΔΔCT^ method.

### Electrophysiology

Sodium current was recording in standard whole-cell patch clamp. The perfusion medium for recording *I*_Na_ was composed, in mmol/L: NaCl 25, CsCl 108.5, HEPES 10, glucose 10, MgCl_2_ 2.5, CoCl_2_ 2.5, CaCl_2_ 0.5 and supplemented with 1 µmol/L of ryanodine, pH adjusted to 7.4 with CsOH. The intrapipette conducting medium was composed, in mmol/L: NaCl 5, MgCl_2_ 2, CaCl_2_ 1, EGTA 15, HEPES 10, MgATP 4, CsCl 130, pH adjusted to 7.2 with CsOH. For the current-Vm and activation (m_∞_) protocol, currents were elicited by 50ms depolarizing pulses from a holding potential of −120 mV, in 5 or 10 mV increments, between −100 and +60 mV, at a frequency of 0.2 Hz. For the steady-state availability-V_m_ protocol (h_∞_), currents were elicited by a 50ms test pulse to −20 mV following 1s pre-pulses applied in 5 or 10 mv increments, between −140 mV and −20 mV, from a holding potential of −120 mV, at 0.2 Hz.

### Statistics

Data were tested for normality using the D’Agostino and Pearson normality test. Statistical analysis was performed using Student’s t test or ANOVA followed by post-hoc Student-Newman-Keuls test on raw data. Results are given as means ± SEM and P values of less than 0.05 were considered significant. Legends: ns, not significant; * P<0.05; ** P<0.01 and *** P<0.001.

## Non-standard Abbreviations and Acronyms

MAGUK: Membrane Associated GUanylate Kinase
CASK: Calcium/CAlmodulin-dependent Serine Kinase
TIRFm: Total Internal Reflection Fluorescence microscopy

## ACKNOWLEDGMENTS

We thank Dr. Rachel Peat for careful reading of the manuscript and helpful comments. This work was supported by the European Union (EUTRAF-261057; EB and SNH), and by the AFM-Telethon doctoral fellowship (21324; AB). The authors declare no competing financial interests.

## AUTHOR CONTRIBUTIONS

Conceptualization: E. Balse, C.A. Eichel, and S. Hatem. Investigation: A. Beuriot, C.A. Eichel, G. Dilanian, F. Louault, D. Melgari, N. Doisne and E. Balse. Formal analysis: A. Beuriot, A. Coulombe and E. Balse. Funding acquisition: E. Balse and S. Hatem. Supervision: E. Balse and S. Hatem. Writing: E. Balse and A. Beuriot.

