## Supplemental Figures for "Distinct CASK domains control cardiac sodium channel membrane expression and focal adhesion anchoring"

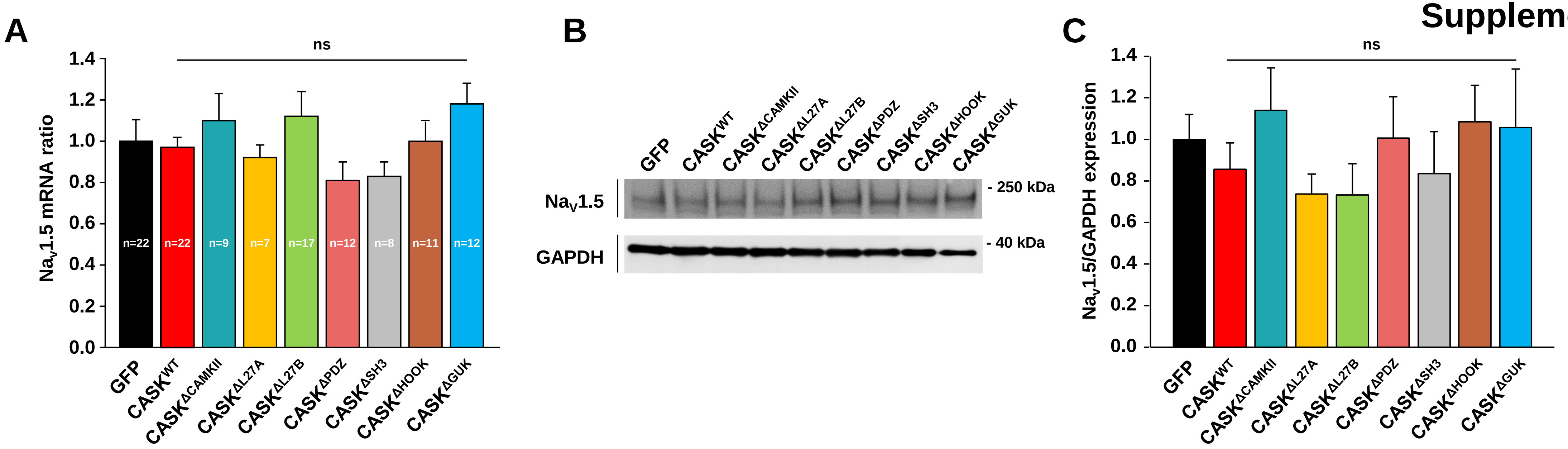


**Supplemental Figure 1. CASK does not regulate translational or transcriptional levels of Na$_V$1.5. (A)** mRNA levels of Na$_V$1.5 in cardiomyocytes 3 days after transduction with the different CASK constructs quantified by RT-qPCR and normalized to GFP control. Legend: ns, not significant, N=3-13 independent cell cultures. **(B)** Representative western blot of Na$_V$1.5 expression after transduction with either GFP, CASK$^{WT}$, or CASK$^{\Delta X}$. **(C)** Corresponding histograms showing the expression levels of Na$_V$1.5 normalized to GAPDH in cardiomyocytes transduced with either GFP, CASK$^{WT}$ or CASK$^{\Delta X}$. Legend: ns, not significant; N=7 independent cell cultures.

### Supplemental Table 1

**A**

**Peak current density at - 20 mV**

| | Mean raw data (pA/pF) | Current density (% from GFP) | Cell capacitance (pF) |
|---|---|---|---|
| GFP | - 70.4 ± 2.9 (n=25) | 100 ± 4.1 (n=25) | 53.1 ± 2.4 (n=25) |
| $CASK^{WT}$ | - 39.6 ± 1.9 (n=36) *** | 56.3 ± 2.7 (n=36) *** | 48.8 ± 2.4 (n=36) $^{ns}$ |
| $CASK^{\Delta CAMKII}$ | - 36.2 ± 3.7 (n=14) *** | 51.4 ± 5.2 (n=14) *** | 47.6 ± 2.5 (n=14) $^{ns}$ |
| $CASK^{\Delta L27A}$ | - 42.4 ± 2.5 (n=7) *** | 60.3 ± 3.5 (n=7) *** | 46.0 ± 3.4 (n=7) $^{ns}$ |
| $CASK^{\Delta L27B}$ | - 65.9 ± 5.3 (n=8) $^{ns}$ | 93.7 ± 7.6 (n=8) $^{ns}$ | 54.5 ± 6.0 (n=8) $^{ns}$ |
| $CASK^{\Delta PDZ}$ | - 38.3 ± 4.3 (n=9) *** | 54.4 ± 6.0 (n=9) *** | 54.2 ± 5.9 (n=9) $^{ns}$ |
| $CASK^{\Delta SH3}$ | - 34.6 ± 4.4 (n=6) *** | 49.1 ± 6.3 (n=6) *** | 44.5 ± 3.8 (n=6) $^{ns}$ |
| $CASK^{\Delta HOOK}$ | - 40.0 ± 4.9 (n=7) *** | 56.9 ± 7.0 (n=7) *** | 49.2 ± 6.5 (n=7) $^{ns}$ |
| $CASK^{\Delta GUK}$ | - 58.9 ± 6.2 (n=10) $^{ns}$ | 83.7 ± 8.9 (n=10) $^{ns}$ | 53.5 ± 3.5 (n=10) $^{ns}$ |

**B**

**Activation properties**

| | Membrane potential at half activation $V_{0.5}$ (mV) | Slope factor k (mV) |
|---|---|---|
| GFP | -39.9 ± 0.7 (n=28) | 6.8 ± 0.2 (n=28) |
| $CASK^{WT}$ | -38.4 ± 0.5 (n=40) $^{ns}$ | 7.2 ± 0.1 (n=40) $^{ns}$ |
| $CASK^{\Delta CAMKII}$ | -41.0 ± 0.3 (n=14) $^{ns}$ | 7.4 ± 0.3 (n=14) $^{ns}$ |
| $CASK^{\Delta L27A}$ | -36.5 ± 2.2 (n=8) $^{ns}$ | 7.3 ± 0.3 (n=8) $^{ns}$ |
| $CASK^{\Delta L27B}$ | -40.4 ± 1.4 (n=12) $^{ns}$ | 7.5 ± 0.3 (n=12) $^{ns}$ |
| $CASK^{\Delta PDZ}$ | -37.4 ± 0.2 (n=9) $^{ns}$ | 8.3 ± 0.9 (n=9) $^{**}$ |
| $CASK^{\Delta SH3}$ | -38.7 ± 2.6 (n=7) $^{ns}$ | 6.8 ± 0.2 (n=7) $^{ns}$ |
| $CASK^{\Delta HOOK}$ | -39.4 ± 1.5 (n=7) $^{ns}$ | 7.39 ± 0.5 (n=7) $^{ns}$ |
| $CASK^{\Delta GUK}$ | -40.7 ± 0.5 (n=10) $^{ns}$ | 6.7 ± 0.2 (n=10) $^{ns}$ |

**C**

**Inactivation properties**

| | Membrane potential at half inactivation $V_{0.5}$ (mV) | Slope factor k (mV) |
|---|---|---|
| GFP | -92.6 ± 0.8 (n=25) | -6.8 ± 0.2 (n=25) |
| $CASK^{WT}$ | -89.6 ± 0.9 (n=27) $^{ns}$ | -6.2 ± 0.2 (n=27) $^{ns}$ |
| $CASK^{\Delta CAMKII}$ | -93.2 ± 1.5 (n=12) $^{ns}$ | -5.9 ± 0.4 (n=12) $^{ns}$ |
| $CASK^{\Delta L27A}$ | -90.0 ± 2.1 (n=12) $^{ns}$ | -6.3 ± 0.2 (n=12) $^{ns}$ |
| $CASK^{\Delta L27B}$ | -89.7 ± 3.2 (n=5) $^{ns}$ | -6.6 ± 0.4 (n=5) $^{ns}$ |
| $CASK^{\Delta PDZ}$ | -95.7 ± 2.7 (n=5) $^{ns}$ | -6.0 ± 0.3 (n=5) $^{ns}$ |
| $CASK^{\Delta SH3}$ | -92.8 ± 3.0 (n=7) $^{ns}$ | -6.0 ± 0.5 (n=7) $^{ns}$ |
| $CASK^{\Delta HOOK}$ | -95.7 ± 1.2 (n=10) $^{ns}$ | -7.5 ± 0.4 (n=10 ) $^{ns}$ |
| $CASK^{\Delta GUK}$ | -95.5 ± 1.2 (n=13) $^{ns}$ | -6.5 ± 0.2 (n=13) $^{ns}$ |

**Supplemental Table 1. Summary table of electrophysiological parameters of cardiac sodium current recorded in adult rat cardiomyocytes 3 days after transduction with adenoviral constructs. (A)** Mean current density of $I_{Na}$ recorded at -20 mV and cell capacitances. **(B)** Activation parameters. **(C)** Steady-state inactivation parameters. Legend: ns, not significant; ** P<0.01; *** P<0.001; n=number of cells; N=3-16 independent cell cultures.
